# FGFR4 phosphorylates MST1 to confer breast cancer cells resistance to MST1/2-dependent apoptosis

**DOI:** 10.1101/431783

**Authors:** S. Pauliina Turunen, Pernilla von Nandelstadh, Tiina Öhman, Erika Gucciardo, Beatriz Martins, Ville Rantanen, Huini Li, Katrin Höpfner, Markku Varjosalo, Kaisa Lehti

**Affiliations:** Department of Microbiology, Tumor and Cell Biology (MTC), Karolinska Institutet, Stockholm, SE-171 77, Sweden; Research Programs Unit, Genome-Scale Biology, Medicum, Faculty of Medicine, University of Helsinki and Helsinki University Hospital, Helsinki, FI-00014, Finland; Institute of Biotechnology, Helsinki Institute of Life Science, University of Helsinki, Helsinki, FI-00014, Finland

## Abstract

Cancer cells balance with the equilibrium of cell death and growth to expand and metastasize. The activity of mammalian sterile20-like kinases MST1/2 has been linked to apoptosis and tumor suppression via YAP/Hippo pathway dependent and independent mechanisms. With a kinase substrate screen we identified here MST1 and MST2 among the top substrates for fibroblast growth factor receptor 4 (FGFR4). In COS-1 cells, MST1 was phosphorylated at Y433 residue in an FGFR4 kinase activity-dependent manner, as assessed by mass spectrometry. Blockade of this phosphorylation by Y433F mutation induced MST1 activation, as reflected by increased autophosphorylation at T183 in FGFR4 overexpressing MDA-MB-231 cells. Importantly, the specific short-term inhibition or knockdown of FGFR4 also led to MST1/2 activation in conjunction with induction of MST1/2-dependent apoptosis in an endogenous model of HER2^+^ breast cancer cells. Moreover, FGFR4 knockdown increased the level of active nuclear MST1 coincidentally with cell polarization and membrane-association of YAP in three-dimensional breast cancer cell spheres. Consistently, FGFR4 overexpression correlated with reduced Hippo pathway-mediated, nuclear translocation-inhibiting YAP phosphorylation, and abysmal HER2^+^ breast carcinoma patient outcome in TCGA cohort. Our results reveal a novel mechanism for FGFR4 oncogenic activity via suppression of the stress-associated MST1/2-dependent apoptosis machinery in the tumor cells with prominent HER/ERBB signaling driven proliferation.

## Introduction

Cancer cells rely on oncogenic signaling by receptor tyrosine kinases (RTKs) to drive tumor initiation and progression [1]. When altered upon tumor evolution, RTKs contribute to the development of resistance towards initially effective anti-cancer treatments. Due to inhibitable enzyme activity and cell surface localization, RTKs thus serve as attractive targets for therapy. Fibroblast growth factor receptors (FGFRs) are RTKs that trigger a series of intracellular signaling cascades that can control key cellular processes including survival, proliferation, differentiation, and migration/invasion, as well as angiogenesis – each abnormally regulated in cancer [2]. Four homologous FGFRs are expressed in humans [2]. Unlike the other family members, FGFR4 is dispensable for normal development in mice [3]. This coupled with specific FGFR4 induction in certain cancers, as well as structural differences and drug selectivity relative to other FGFRs, supports the possibility of effective FGFR4 targeting as a therapeutic intervention [4–9].

Despite advances in diagnosis and treatment, subsets of breast cancer remain challenging to cure accounting for an estimated 15% of all cancer deaths in women (World Cancer Report, WHO 2014). Like in other cancers, the poor prognosis is partially attributed to therapy resistance and anti-apoptosis responses of the cancer cells [10]. FGFR4 overexpression and gene alterations, including G388R single nucleotide polymorphism (SNP), have been associated with cancer invasion, drug resistance and poor prognosis [11–13]. Among breast cancers, FGFR4 is overexpressed in a significant proportion of especially HER2-enriched tumors [4], where it has been linked to tumor cell growth and apoptosis resistance [4,5,14]. Despite these results, the molecular mechanisms how FGFR4 confers the aggressive cancer cell behavior remain incompletely understood.

In this study, we systematically screened for FGFR4 substrates using an *in vitro* human kinase substrate protein microarray. Unexpectedly, Hippo tumor suppressor pathway components, including the serine/threonine kinases MST1/2 (mammalian sterile20-like kinases), were among the most prominent tyrosine-phosphorylated substrates for FGFR4. Cytoplasmic MST1/2 comprises the core kinase complex of the mammalian Hippo pathway, which activation ultimately leads to serine phosphorylation-dependent cytoplasmic retention and inactivation of the oncogenic transcriptional regulators, YES-associated protein (YAP1) and transcriptional co-activator with PDZ-binding motif (TAZ) [15,16]. Upon cell stress and apoptosis, caspase-3 cleavage in turn removes the inhibitory C-terminal domains of MST1/2 to induce transport of the activated N-terminal MST1/2 into the nucleus [17]. Although results from overexpression experiments have shown that nuclear MST1/2 can promote apoptosis [18–21], and reduced MST1/2 serine/threonine kinase activity has been linked to poor cancer prognosis [22–25], the functional contribution of endogenous MST1/2 in physiological or pathological apoptotic processes have remained elusive [17]. Our screening results led us to investigate in more detail the putative direct functional interaction of FGFR4 and MST1/2. Altogether, these results establish a unique mechanism of oncogenic signaling – the dominant FGFR4-mediated attenuation of MST1/2-mediated, stress-associated apoptosis in HER2^+^ breast cancer.

## Materials and Methods

### Kinase Substrate Identification Array

For *in vitro* FGFR4 substrate identification, the protein microarray containing 9483 human recombinant proteins (Protoarray human protein microarrays version 5.0; Invitrogen), was blocked and incubated with the recombinant kinase domain of FGFR4 (50 nM; Invitrogen) in the presence of [γ-33P] ATP. The array was washed to remove unbound γ-33P, and exposed to X-ray film. The acquired array image was analyzed using ProtoArray Prospector software (Invitrogen). The raw data were subjected to background subtraction, signal scatter compensation, and outlier detection. The Z factor cutoff value was set at ≥0.4. Phosphorylated proteins with a Z score > 0.25 were considered as potential substrates.

### Cell Culture

Human ZR-75.1, MCF7, BT474, T47D, MDA-MB-453, Hs578T, BT549, MDA-MB-231 (American Type Culture Collection; ATCC) and SUM159 [26] breast carcinoma cells, and COS-1 cells (ATCC) were cultured according to manufacturer’s instructions. The MycoAlertPlus kit (LT07-705, Lonza) was used for confirming the cell cultures negative for mycoplasma.

### Antibodies, inhibitors and growth factors

The antibodies used were as follows: mouse monoclonal antibodies against ERBB2/HER2 (MA5-13105, Thermo Fisher Scientific for IF, and NCL-L-CB11; Leica Biosystems), FGFR4 (sc-136988, Santa Cruz Biotechnology), FLAG (9A3; 8146, Cell Signaling Technology), glyceraldehyde-3-phosphate dehydrogenase (GAPDH; G8795; Sigma-Aldrich), MST1 (sc-100449, Santa Cruz Biotechnology, for IF), phospho-tyrosine (05-321; EMD Millipore Corporation), YAP (sc-101199, Santa Cruz Biotechnology), rabbit monoclonal antibodies against MOB1 (13730S), phospho-MOB1 (Thr35) (D2F10; 8699) and phospho-tyrosine (8954, MultiMab) all form Cell Signaling Technology, and rabbit polyclonal antibodies against FGFR4 (sc-124; Santa Cruz), phospho-FRS2-α (Tyr196; 3864), MST1 (3682), MST2 (3952), phospho-p44/42 MAPK (phospho-Erk1/2) (9101), phospho-AKT (Ser473; 9271), phospho-MST1 (Thr183)/MST2(Thr180); 3681), phospho-YAP (Ser127; 4911) (Cell Signaling Technology), V5-tag (ab9116; Abcam), ZO-1 (61-7300; Zymed, Invitrogen), and horseradish peroxidase-conjugated secondary antibodies (P044701 and P044801, Dako) for enhanced chemiluminescence detection of immunoblots. Alexa Fluor conjugated secondary antibodies and phalloidin (Life Technologies) were used for immunofluorescence. FGFR4 inhibitor BLU9931 (S7819) was purchased from Selleckchem. Recombinant human fibroblast growth factor 1 (GF002) and fibroblast growth factor 2 (GF003) were purchased from Merck.

### cDNA constructs, small interfering RNAs, and short hairpin RNAs

Flag epitope (N-terminal)-tagged MST1 and MST2 in the p3FLAG-CMV-10wt (Sigma-Aldrich) expression vectors have been described previously [27]. MST1 Y433F mutant was generated by site-directed mutagenesis. FGFR4 and its kinase activity-deficient mutant (K503M) were cloned into pcDNA3.1/V5-His vector, and the expression vector for FGFR4 G388 variant was generated from R388 variant by site-directed mutagenesis [28,29]. siRNAs used were SMARTpool siGENOME (Dharmacon, GE Healthcare) against human STK3 (MST2; L-004874), STK4 (MST1; L-004157) and nontargeting control siRNA (D-001206-14-10), as well as siRNAs against FGFR4 (ON-TARGETplus siRNA pool, L-003134 Dharmacon, GE Healthcare; or SI02665306 (siFGFR4_6) and SI00031360 (siFGFR4_2), Qiagen).

Short hairpin RNA (shRNA) targeted against FGFR4 (TRCN0000000628, TRCN0000010531 and TRCN0000199510; Open Biosystems, GE Healthcare) or nontargeting scrambled shRNA were used. The packaging plasmid (pCMVdr8.74), envelope plasmid (pMD2-VSVG), and FGFR4 or scrambled shRNA in pLKO.1 vector were co-transfected into 293FT producer cells using Lipofectamine 2000 reagent (Invitrogen). Complete breast carcinoma cell growth medium was changed on 293FT cells 24 hours after transfection. The viral supernatants were collected after 48 hours, passed through a 0.45-μm filter, and incubated with human breast carcinoma cells. After 16 hours of infection, the supernatants were replaced with complete media followed by puromycin (2 μg/ml) selection of the transduced cells [30].

### Cell transfections, sphere preparation and treatments

The cells were transfected with expression vectors using FuGENE HD (Promega, Madison, WI) and siRNAs using Lipofectamine 2000 (Invitrogen) or Interferin (Polyplus-transfection SA). Knockdown efficiencies were analyzed by Western blotting after 72 hours. For 2D immunofluorescence, cells were seeded on monomeric collagen coated (type I, 50 μg/ml, Sigma-Aldrich) coverslips, fixed in 4% paraformaldehyde (pH 7.5), and stained as previously described [31], and mounted in Vectashield with 4’,6-diamidino-2-phenylindole (DAPI; Vector Laboratories).

Spheres of 24 000 cells were allowed to form under nonadherent conditions in agarose-coated 96-well plates in cell culture media supplemented with 0.2 mg/ml Matrigel (growth factor reduced; Corning) and cultured for 5–8 days [32]. The spheres were cultured in the indicated serum concentrations, or serum-starved for 16 hours before inhibitor treatment, followed by lysis with RIPA buffer. For immunofluorescence, the spheres were fixed in 4% paraformaldehyde (pH 7.5), post-fixed in ice-cold acetone-methanol (1:1) solution, incubated in blocking buffer (5% BSA, 0.1% Triton X-100 in PBS) and stained for MST1. Nuclei were visualized with Hoechst 33342 stain (Thermo Fisher Scientific). The spheres were mounted on glass slides in Vectashield (Vector Laboratories). At least two different cell lines and two independent cultures per stable lentiviral shRNA transduction were analyzed for MST1.

### Immunoprecipitation and immunoblotting

Immunoprecipitation and immunoblotting were performed as described previously [33]. Cells were lysed with RIPA buffer (50 mM Tris-HCl pH 7.4, 150mM NaCl, 1% Igepal CA-630, 0.5% sodiumdeoxycholate) containing Complete protease inhibitor cocktail (Roche), 2 mM Na_3_VO_4_, and 2 mM NaF and cleared by centifugation. For immunoprecipitation from the soluble cell lysates, the samples were further diluted 1:3 with Triton lysis buffer (50 mM Tris-HCl pH 8.0, 150mM NaCl, 1% Triton X-100, 5 mM CaCl_2_, 0.02% NaN_3_) containing Complete protease inhibitor cocktail, 2 mM Na_3_VO_4_, and 2 mM NaF and pre-cleared with protein A Sepharose. MST1 or MST2 were immunoprecipitated from cleared supernatants with EZview Red anti-FLAG M2 Affinity Gel (Sigma-Aldrich) for 4 hours at 4°C. After washing with Triton lysis buffer, bound proteins were eluted with reducing SDS-PAGE sample buffer (0.12 M Tris-HCl pH 6.8, 0.02% bromophenol blue, 4% SDS, 50% glycerol). Equivalent quantities of protein were separated by SDS-PAGE and transferred to nitroclellulose membranes for subsequent detection with primary antibodies and matched secondary antibodies conjugated to HRP.

### Mass spectrometry analysis of phosphorylation sites

All buffers of immunoprecipitation and elution procedures were supplemented with Complete protease inhibitor cocktail and PhosSTOP phosphatase inhibitor tablets (Roche). After the general immunoprecipitation washes, the anti-FLAG affinity gel was washed with pre-urea wash buffer (50 mM Tris pH 8.5, 1 mM EGTA, 75 mM KCl) before elution with urea buffer (6M Urea, 20 mM Tris pH 7.5, 100 mM NaCl). Elution cycles were repeated three times for 30 minutes each at RT with agitation. Proteins in eluates were reduced with (tris(2-carboxyethyl)phosphine), alkylated with iodoacetamide, and trypsin digested with Sequencing Grade Modified Trypsin (Promega). Phosphopeptide enrichment was performed using immobilized metal ion affinity chromatography with titanium (IV) ion (Ti4+-IMAC). IMAC material was prepared and used essentially as described [34]. The LC-MS/MS analysis was performed using a Q Exactive ESI-quadrupole-orbitrap mass spectrometer coupled to an EASY-nLC 1000 nanoflow LC (Thermo Fisher Scientific), using Xcalibur version 3.1.66.10 (Thermo Fisher Scientific). The phoshopeptide sample was loaded from an autosampler into a C18-packed precolumn (Acclaim PepMap™100 100 μm x 2 cm, 3 μm, 100 Å, Thermo Fisher Scientific) in buffer A (1 % acetonitrile (ACN), 0.1 %, formic acid (FA)). Peptides were transferred onward to a C18-packed analytical column (Acclaim PepMap™100 75 μm × 15 cm, 2 μm, 100 Å, Thermo Fisher Scientific) and separated with 60-minute linear gradient from 5 to 35 % of buffer B (98 % ACN, 0.1 % FA) at the flow rate of 300 nl/minute. The mass spectrometry analysis was performed in data-dependent acquisition in positive ion mode. MS spectra were acquired from m/z 300 to m/z 2000 with a resolution of 70,000 with Full AGC target value of 3,000,000 ions, and a maximal injection time of 120 ms, in profile mode. The 10 most abundant ions with charge states from 2+ to 7+ were selected for subsequent fragmentation (higher energy collisional dissociation, HCD) and MS/MS spectra were acquired with a resolution of 17,500 with AGC target value of 5000, a maximal injection time of 120 ms, and the lowest mass fixed at m/z 120, in centroid mode. Dynamic exclusion duration was 30 s. Raw data files were analyzed with the Proteome Discoverer software version 1.3 (Thermo Fisher Scientific) connected a Sequest search engine version 28.0 (Thermo Fisher Scientific) against the human component of the Uniprot Database (version 07_2017). Carbamidomethylation (+57.021 Da) of cysteine residues was used as static modification. Phosphorylation of Ser/Thr/Tyr (+79.966 Da) and oxidation (+15.994 Da) of methionine was used as dynamic modification. Precursor mass tolerance and fragment mass tolerance were set to less than 15 ppm and 0.05 Da, respectively. A maximum of two missed cleavages was allowed. The software phosphoRS [35] was used to calculate the invidual site probabilities for phosphorylated peptides.

### Flow cytometry analysis of apoptosis

The proportions of apoptotic cells were determined by flow cytometry for annexin V and propidium iodide binding [36]. Stable shScr or shFGFR4 transduced MDA-MB-453 cells were transfected with siRNAs. After 72 hours the cells, including floating cells in the medium, were collected by trypsinization, washed and labeled with annexin V-Alexa Fluor488 (A13201; Life Technologies) diluted into binding buffer (10 mM HEPES – 140 mM NaCl – 2.5 mM CaCl_2_) for 15 min at RT. Cells were suspended into binding buffer, stained with 1 μg/ml propidium iodide (P3566, Life Technologies), and analyzed with BD Accuri C6 flow cytometer. Gating and data analysis were performed using FlowJo v10.1 software (Tree Star Inc). Geometric mean fluorescence of annexin V binding (FL1-A) alone or combined with propidium iodide binding (FL3-A) was quantified.

### Imaging and image quantification

Fluorescence images were obtained using an AxioImager.Z2 upright epifluorescence microscope with Plan-Apochromat 20×/0.8 NA dry or 40×/1.4 NA oil objective. In addition, an LSM 780 confocal microscope with Plan-Neofluar 40×, 1.3 NA oil objective was used (all from Carl Zeiss). Brightness and contrast were linearly adjusted using ZEN 2012 (blue edition; Carl Zeiss) or Photo-Paint X7 (Corel). Single optical sections or a combination of two serial optical sections were used for image display.

### Image quantifications of Western blotting were performed by processing all obtained micrographs with ImageJ software

The MST1 and YAP-stained sphere images were analyzed using Anima [37]. Cell nuclei were detected from DAPI channel by first normalizing images with adaptive histogram equalization, and then using the Shape filtering segmentation method found in Anima. The MST1 or YAP signal intensity values were measured from the nucleus area and from a ring around the nucleus representing the cytoplasm. The ring width was determined as 50% of the radius of the nucleus. The nucleus/cytoplasm intensity ratio was calculated by dividing nuclear intensity with cytoplasmic intensity for each nucleus, and then calculating a median value for each image.

### Statistics

All numerical values represent mean ± SD or SEM as indicated in figure legends. Statistical significance was determined using two-tailed Student’s t tests for analysis of apoptosis and Western blot quantifications. The Kolmogorov-Smirnov test was used for computerized image analysis data on MST1 and YAP-stained sphere images.

## Results

### FGFR4 tyrosine-phosphorylates Hippo pathway proteins *in vitro* and in COS-1 cells

To systematically screen for FGFR4 substrates that serve as downstream oncogenic effectors, we used recombinant human FGFR4 kinase domain to assess *in vitro* phosphorylation of 9483 human recombinant proteins (Fig. 1A). Unexpectedly, the top five FGFR4 substrates ranked by the Z-score included four Hippo tumor suppressor pathway proteins; MST2 *(STK3),* protein kinase C iota *(PRKCI),* casein kinase I delta *(CSNK1D),* and MST1 *(STK4)* (Fig. 1B; STK3 and PRKCI identified as two splice variants), suggesting that FGFR4 tyrosine kinase activity can directly affect Hippo serine/threonine kinase signaling (Fig. 1B). To validate the FGFR4 activity in cells, the core Hippo kinases MST1/2 were immunoprecipitated from COS-1 cells after transfection of MST1 and MST2 alone or in combination with FGFR4 G388 (G) or the cancer-associated R388 (R) SNP variant. Notably, both variants induced tyrosine phosphorylation of MST1 and MST2, establishing these Hippo kinases as novel FGFR4 substrates (Fig. 1C). Importantly, the FGFR4 (R)-mediated MST1/2 tyrosine phoshorylation was not detected after co-transfection with kinase activity-dead FGFR4 (R) KD (Fig. 1D).

**Figure 1.**
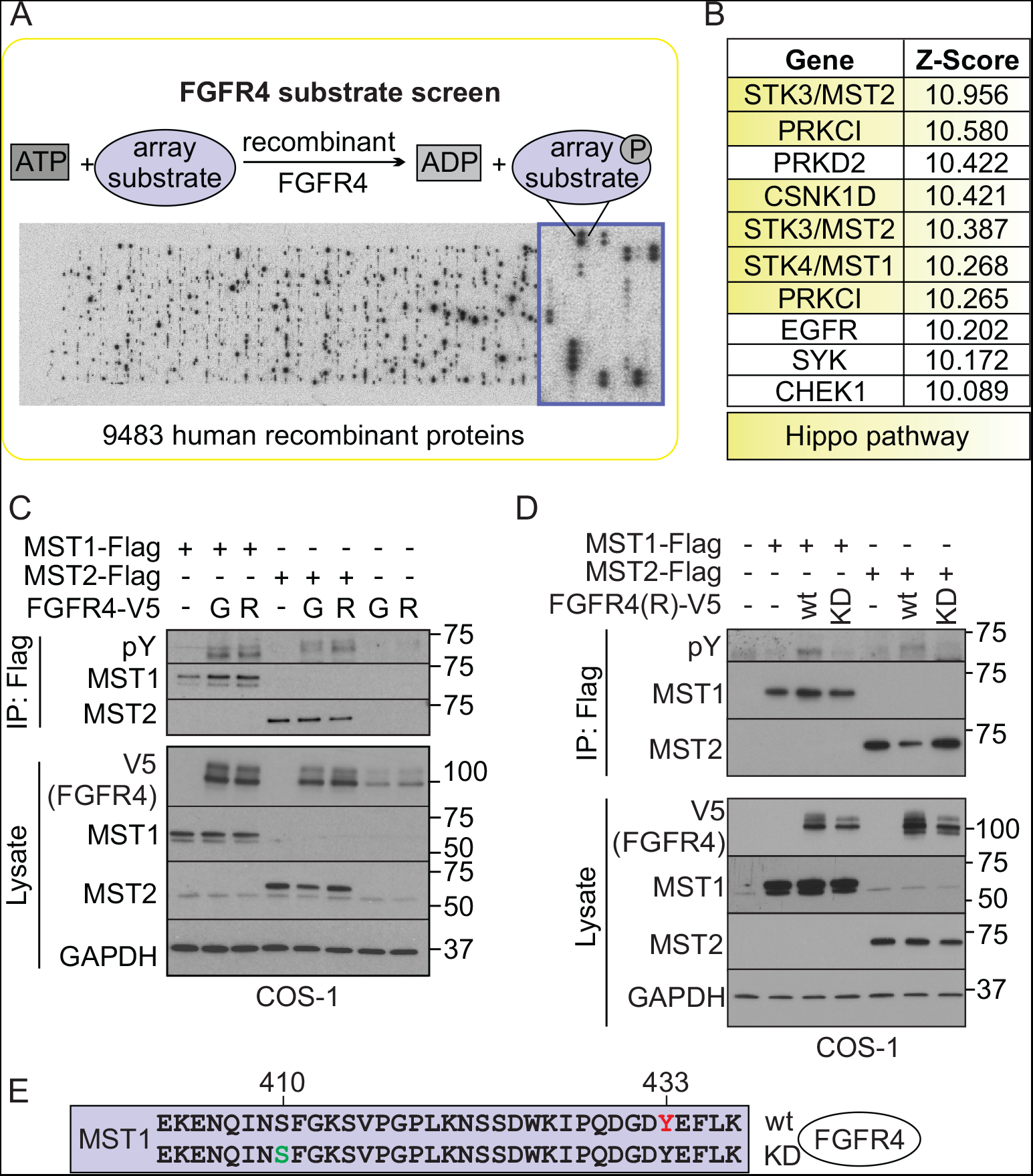
FGFR4 substrate screen identifies tyrosine phosphorylated Hippo pathway proteins including MST1/2. **(A)** Scheme of the substrate screen with recombinant FGFR4 kinase domain. (**B)** Top 10 FGFR4 substrates ranked by the Z-score include Hippo pathway-associated proteins (yellow). (**C, D)** MST1/2 are tyrosine phosphorylated by FGFR4 in COS-1 cells. Flag-tagged MST1/2 were immunoprecipitated after transfection of MST1 and MST2 alone or in combination with FGFR4 G388 (G), or R388 (R) kinase (wt) or kinase-dead (KD) variants, and detected by immunoblotting. **(E)** MST1 immunoprecipitates from COS-1 cells co-transfected with FGFR4 (R)-wt or FGFR4 (R)- KD (See Fig. S1A) were trypsin digested and subjected to phoshopeptide enrichment prior to LC-MS/MS analysis that identified phosphorylated Y433 (red) on MST1 only with FGFR4 (R)-wt, and phosphorylated S410 (green) only with FGFR4 (R)-KD.

To identify the FGFR4 phosphorylated tyrosine residue(s), the immunoprecipitated MST1 was subjected to mass spectrometry (Fig. S1A). Notably, MST1 derived from the FGFR4 expressing COS-1 cells was phosphorylated at Y433, whereas MST1 from the cells with inactive FGFR4 KD lacked tyrosine phosphorylated residues (Fig. 1E, Table 1). Additionally, an uncharacterized MST1 phosphorylation at S410 near the pY433-site was detected in the absence of FGFR4 kinase activity (Fig. 1E, Table 1), whereas phosphorylation of MST1 S320 and T177 appeared independently of FGFR4 activity (Table 1).

**Table 1.** List of MST1 phoshopeptides identified by mass spectrometry.

| **Co-transfected kinase** | **Protein** | **Phospho-residue** | **Peptide Sequence** | **Peptide FDR** | **phosphoRS Site probabilities (%)** |
| --- | --- | --- | --- | --- | --- |
| FGFR4/R-wt | MST1_Q13043 | Y(433) | IPQDGD<u>Y</u>EFLK | <0.01 | 100 |
|  | MST1_Q13043 | S(320) | EVDQDDEEN<u>S</u>EEDEMDSGTMVR | <0.01 | 100 |
|  | MST1_Q13043 | S(410) | EKENQIN<u>S</u>FGK | N/A | 0 |
|  | MST1_Q13043 | T(177) | LADFGVAGQLTD<u>T</u>MAK | <0.01 | 99.9 |
| FGFR4/R-KD | MST1_Q13043 | Y(433) | IPQDGD<u>Y</u>EFLK | N/A | 0 |
|  | MST1_Q13043 | S(320) | EVDQDDEEN<u>S</u>EEDEMDSGTMVR | <0.01 | 100 |
|  | MST1_Q13043 | S(410) | EKENQIN<u>S</u>FGK | <0.01 | 100 |
|  | MST1_Q13043 | T(177) | LADFGVAGQLTD<u>T</u>MAK | <0.01 | 100 |

### FGFR4 is overexpressed in HER2^+^ breast cancer cells and correlates with adverse outcome in HER2-enriched breast cancer patients

To investigate the clinical relevance of the identified novel FGFR4 activity, and to establish relevant cell models, we first analyzed FGFR4 expression using TCGA by cBioPortal for Cancer Genomics [38,39]. In breast cancer, FGFR4 was overexpressed in 33% of the HER2-enriched tumors (PAM50 classified, n=58 [40], and significantly associated with poorer overall survival as compared to the non-overexpressing group (Fig. S1B; P = 0.044; FGFR4 overexpression in 4% of all breast cancers; TCGA, n=825 [4]). Consistent with the expression in human tumors, MDA-MB-453 (ER^−^, PR^−^, HER2^+^) as well as ZR-75.1 and BT474 (ER^+^, PR^+/-^, HER2^+^) human breast cancer cells expressed FGFR4, whereas both FGFR4 and HER2 were low in MCF7 and T47D cells (ER^+^, PR^+/-^, HER2^−^), and negligible in five triple-negative (ER^−^, PR^−^, HER2^−^) cell lines (Fig. 2A, S1C) [41]. By immunofluorescence, strong intracellular and cell surface FGFR4 was associated with prominent cell surface HER2 in MDA-MB-453 and ZR-75.1 cells, rendering these cells suitable endogenous breast cancer models for FGFR4 function (Fig. 2B).

**Figure 2.**
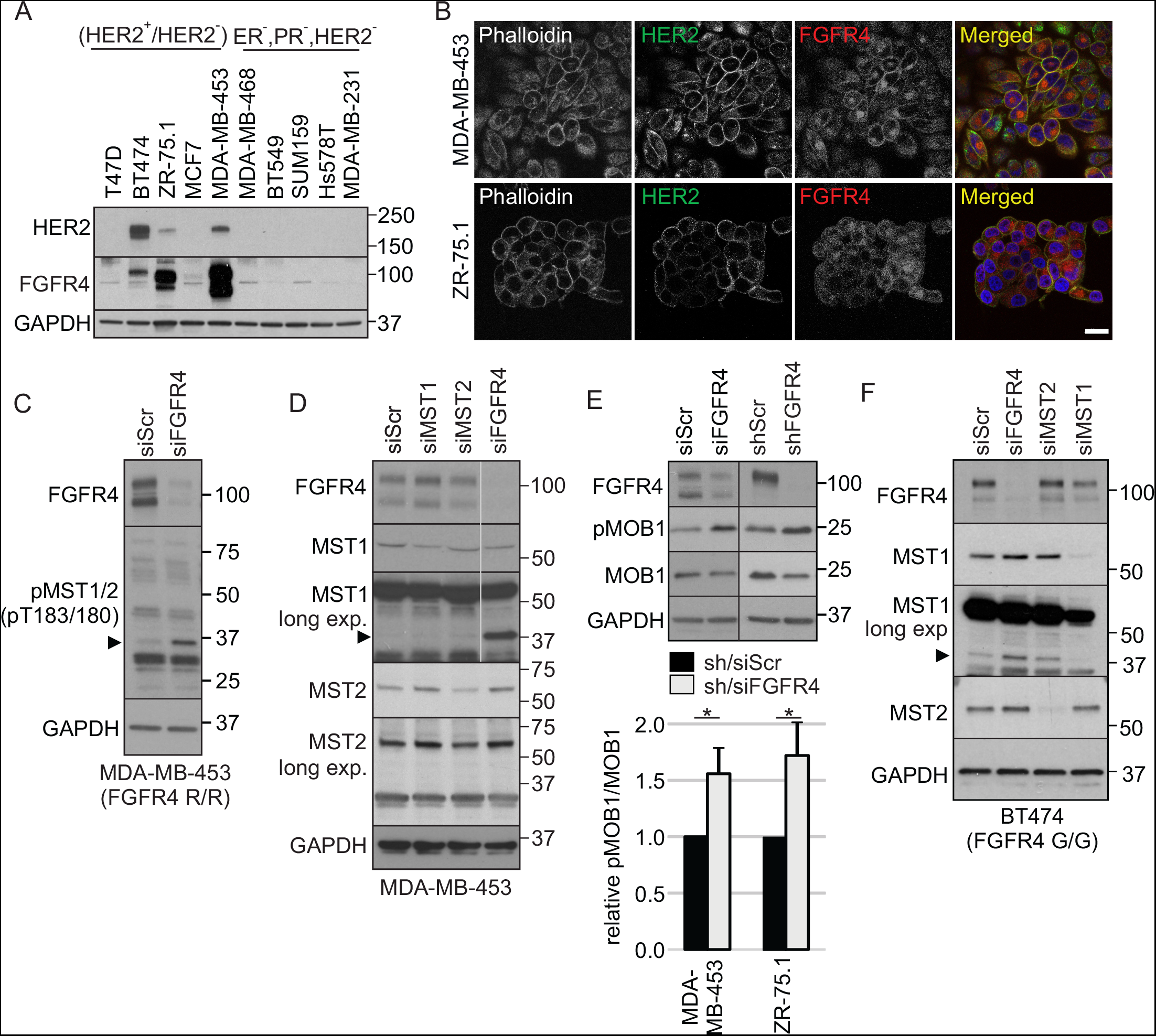
FGFR4 suppresses MST1/2 activation and cleavage in HER2^+^ breast cancer cells. **(A, B)** FGFR4 and HER2 expression in luminal MDA-MB-453, ZR-75.1 and BT474, MCF7 and T47D, and five triple-negative breast cancer cell lines by **(A)** immunoblotting and **(B)** immunofluorescence. Scale bar 20 μm. **(C-D)** MDA-MB-453 cells transfected with indicated siRNAs were subjected to immunoblotting for **(C)** T183/180 phosphorylated MST1/2, and **(D)** MST1 and MST2. Note cleaved ~37 kDa MST1/N in FGFR4 knockdown cells (arrow head in C and D). Thin grey line indicates cropping to leave out irrelevant sample lane; see uncropped immunoblots in Fig. S5. **(E)** MDA-MB-453 and ZR-75.1 cells were transduced with indicated si/shRNAs; upper, indicated immunoblots; lower, quantification of pMOB1/MOB1 ratio, n=3, mean ± SEM; * P<0.05. **(F)** BT474 cells transfected with indicated siRNAs were subjected to immunoblotting for MST1 and MST2.

### FGFR4 inhibits MST1/2 activation and cleavage

To test if endogenous FGFR4 regulates MST1/2 kinases, MDA-MB-453 and ZR-75.1 breast cancer cells were transfected with siRNA pools specific for FGFR4, MST1 and MST2. In control siRNA transfected MDA-MB-453 cells, weak protein bands corresponding to the sizes of full length and cleaved MST1/2 were detected with antibodies specific to the active, pT183/180 autophosphorylated MST1/2 (Fig. 2C, arrow head pointing to cleaved MST1/2). Markedly, FGFR4 knockdown enhanced the cleaved MST1/2 pT183/180 (Fig. 2C, arrowhead). MST1, expressed in control cells mainly as a full-length protein, became detectable as an additional 37 kDa form specifically after FGFR4 knockdown (Fig. 2D; see long exposure for prominent N-terminal fragment similar to the caspase cleaved MST1/N [17,21]. MST2 instead remained essentially unaffected after knockdown of FGFR4 or MST1 (Fig. 2D). Consistently, after stable FGFR4 (R) or (G) overexpression in MDA-MB-231 breast cancer cells, cleaved 37 kDa MST1 was decreased as compared to the control cells (Fig. S2A), and even more efficiently upon FGF1 and FGF2 stimulation as shown in FGFR4 (R) expressing cells (Fig. S2B).

Coincident with the above alterations in MST1/2 activation in MDA-MB-453 cells, knockdown of endogenous FGFR4 resulted in significantly increased threonine (T35) phosphorylation of the MST1/2 substrate MOB1 (Fig. 2E; P<0.05; [42]). FGFR4 silencing by lentiviral shRNAs in ZR-75.1 cells likewise enhanced MOB1 pT35 (Fig. 2E; P<0.05), and MST1/2 pT183/180 (Fig. S2C), consistent with increased MST1/2 activation. This was coupled with enhanced full-length MST1/2, whereas cleaved MST1/2 was less prominent in these cells (Fig. S2C). Since MDA-MB-453 and ZR-75.1 cells both contain the cancer-associated FGFR4 R388 variant (homozygous in MDA-MB-453 and heterozygous in ZR-75.1 by sequencing), we further silenced FGFR4 in BT474 cells homozygous for the alternative G388 variant. FGFR4 silencing likewise increased the full-length MST1/2 as well as cleaved MST1 (Fig. 2F), indicating that distinct from what has been found for FGFR4-induced cell invasion and STAT3 signaling [12,29,43], the FGFR4-mediated MST1 regulation did not correlate with G388R polymorphism.

### FGFR4 counteracts MST1/2-mediated apoptosis

MST1/2 activation by T183/180 autophosphorylation and cleavage is associated with apoptosis [17]. To investigate if FGFR4 alters the pro-apoptotic MST1/2 function, MDA-MB-453 cells were stably transduced with lentiviral control (shScr) and FGFR4 (shFGFR4) shRNAs, followed by transfection of siRNA pools targeting FGFR4, MST1 and MST2. Apoptosis was analysed by measuring annexin V and propidium iodide (PI) binding with flow cytometry using two gating strategies: P1 (smaller) and P2 (larger) cells for early apoptosis detection by annexin V binding (Fig. 3A), and the whole cell population for quantification of combined early and late apoptotic cells by double-positivity (Fig. 3B, C; annexin V + PI). Stable FGFR4 silencing significantly increased apoptosis relative to the control (Fig. S3A; P≤0.02 and Fig. 3A-C). Similarly, FGFR4 siRNAs increased apoptosis in shScr cells (Fig. 3A-C; P<0.001 in C). MST1 or MST2 knockdown did not decrease apoptosis, and even slightly increased apoptosis in shScr cells with high endogenous FGFR4 (Fig. 3A-C). Notably, while FGFR4 siRNAs did not further increase apoptosis in shFGFR4 cells (Fig. 3A-C), the apoptosis induction caused by the stable FGFR4 silencing was rescued close to the low level of shScr cell apoptosis by knockdown of MST1 (Fig. 3A-C, P=0.02 in C) or MST2 (Fig. 3A-C; P=0.003 in C) in shFGFR4 cells. These results indicate that endogenous FGFR4 attenuates the pro-apoptotic MST1/2 activity in MDA-MB-453 cells.

**Figure 3.**
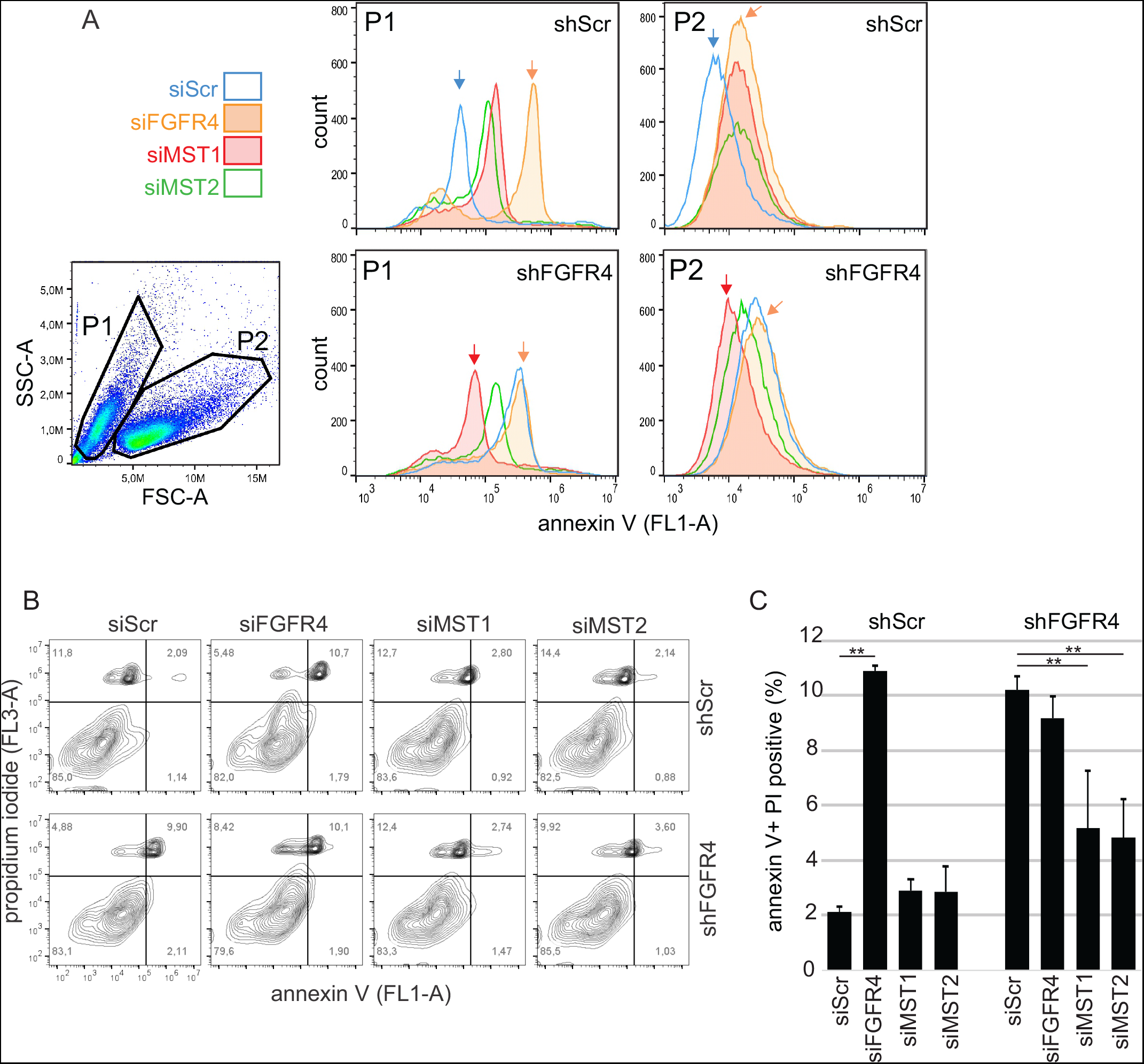
FGFR4 counteracts MST1/2-mediated apoptosis. MDA-MB-453 cells transduced with shScr or shFGFR4 shRNAs were transfected with siRNA pools specific for FGFR4, MST1 or MST2, and analysed for annexin V and propidium iodide (PI) binding by flow cytometry using two different gating strategies for data visualization. **(A)** Gating to populations P1 (smaller) and P2 (larger), and annexin V binding (FL1-A) histograms as a marker for early apoptotic cells. (**B)** Representative contour plots show the annexin V (FL1-A) and PI (FL3-A) double positive cells, indicating both early and late apoptotic stages, in the upper right quadrant (% of total, 100 000 events). **(C)** Quantification of apoptosis based on double-positive (annexin V + PI) cells in the upper right quadrant from a representative experiment; mean ± SD of triplicates, ** P < 0.01. FSC-A; forward scatter, and SSC-A; side scatter.

### FGFR4 silencing increases MST1/2 activation and nuclear translocation in cancer cell spheres

Three-dimensional (3D) cell spheres recapitulate the cell-cell contacts and dimensionality of *in vivo* tumors. To assess the effects of FGFR4 in MST1/2 regulation and apoptosis in 3D, MDA-MB-453 cells were cultured under non-adherent conditions to support sphere formation. The control and FGFR4 knockdown spheres had minor differences in MST1/2 and MOB1 protein and phoshorylation when cultured in complete medium (Fig. 4A; 10% FBS). Notably, FGFR4 depletion in low serum cultures led to increased full-length and cleaved MST1/2, in conjunction with enhanced MST1/2 pT183/180 and MOB1 pT35 in shFGFR4 cells (Fig. 4A; 2% FBS). Treatment of MDA-MB-453 spheres with FGFR4 inhibitor BLU9931 for 15 minutes likewise increased pT183/180 MST1/2 (Fig. 4B). During this short term FGFR4 inhibition FRS2α was diminished, whereas pAKT and pERK remained almost unaffected (Fig. 4B), linking FGFR4 activity directly to MST1 regulation. Moreover, as assessed by immunofluorescence of the MDA-MB-453 and ZR-75.1 spheres, shScr cells had low nuclear and cytoplasmic MST1, whereas FGFR4 silencing increased MST1 specifically in the nuclei, significantly increasing the nucleus/cytoplasm intensity ratio (Fig. 4C, D, S3B). Altogether these results suggest that FGFR4 suppresses MST1/2 activation and MST1 nuclear localization to counteract apoptosis.

**Figure 4.**
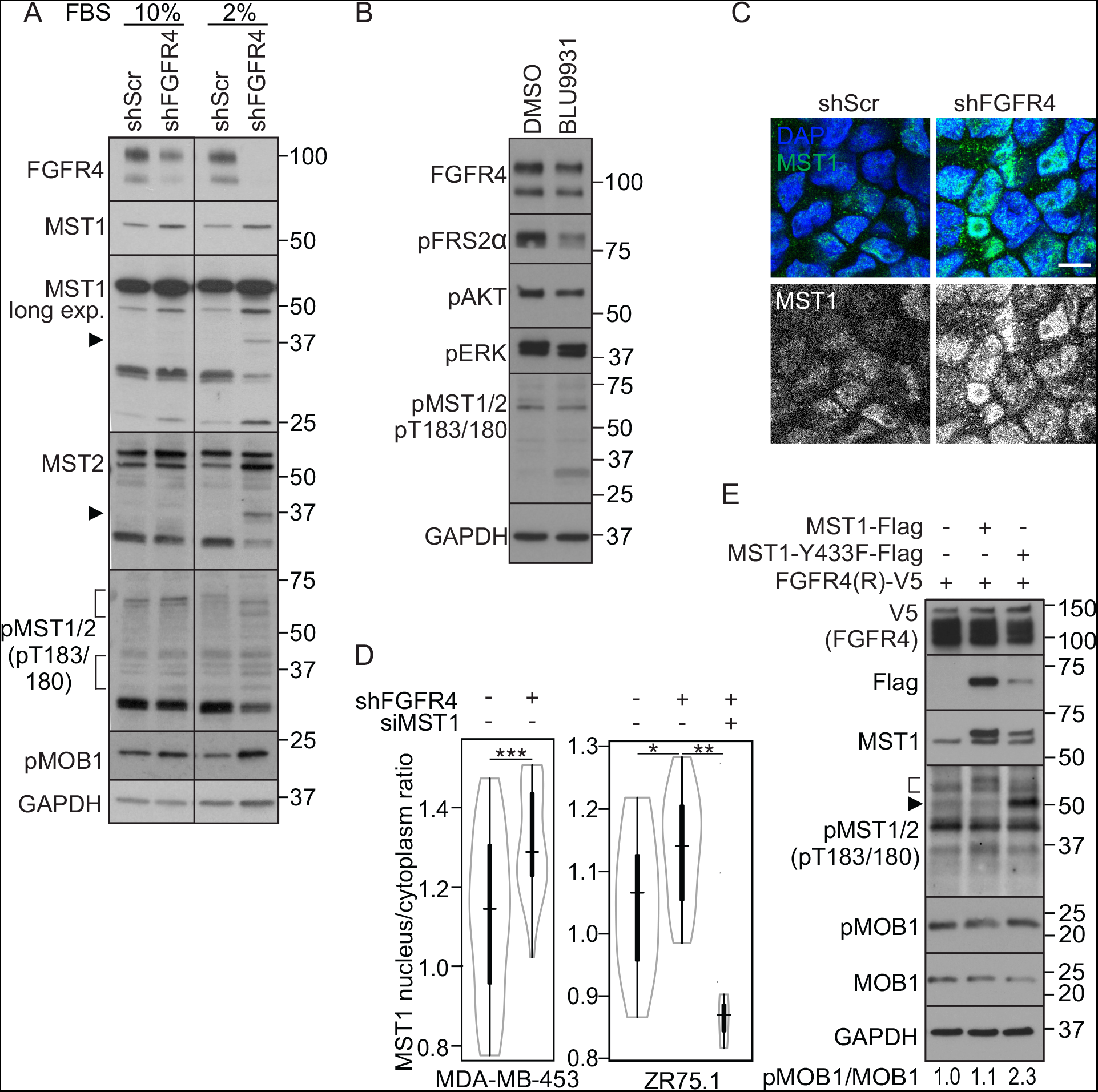
Suppression of MST1/2 activation is directly linked to FGFR4 kinase activity. **(A)** shScr and shFGFR4 MDA-MB-453 cell spheres were cultured under non-adherent conditions (10% or 2% FBS), and subjected to immunoblotting. Arrow heads indicate cleaved N-terminal MST1/2 (2% FBS), brackets highlight the fragments of autoactivated MST1/2. **(B)** MDA-MB-453 cell spheres were treated with 100 nM BLU9931 for 15 minutes, and subjected to immunoblotting. **(C, D)** shScr and shFGFR4 MDA-MB-453 and ZR-75.1 spheres were analyzed for MST1 expression by **(C)** immunofluorescence, and **(D)** MST1 nuclear/cytoplasmic ratio was quantified (n = 4-;6 MDA-MB-453 spheres, ≥6 microscopic fields/sphere; n = 2-3 ZR-75.1 spheres, ≥8 microscopic fields / sphere; mean ± SEM of two independent experiments. Scale bar 10 μm. **(E)** MDA-MB-231 cells cotransfected with FGFR4 (R) and wild-type or phosphosite mutant MST1-Y433F were subjected to immunoblotting as indicated. Bracket and arrow head indicate the activated pT183 MST1 fragments. Ratio of pMOB1/MOB1 is indicated below the immunoblot panel.

### Mutation of the site for FGFR4-mediated tyrosine phosphorylation restores MST1 autoactivation in FGFR4 expressing cells

To directly link the identified FGFR4-dependent tyrosine phosphorylation to MST1/2 regulation, MDA-MB-231 cells were co-transfected with FGFR4 (R) and wild-type MST1 or tyrosine phosphorylation site mutant MST1-Y433F. While weak protein bands corresponding to full-length, autoactivated MST1/2 pT183/180 were detected in cells transfected with MST1 alone (wild-type or mutant, Fig. S3C) or FGFR4 (R) in combination with wild-type MST1, the co-transfection of phosphosite mutant MST1-Y433F resulted in prominent autophosphorylated MST1/2 pT183/180 (Fig. 4E). Phosphorylation of MOB1 was also increased relative to total MOB1 in cells co-transfected with MST1-Y433F and FGFR4 (R) (Fig. 4E). These results further indicate that FGFR4-dependent MST1 phosphorylation at Y433 suppresses MST1/2 activation in the FGFR4 expressing cancer cells.

### FGFR4 overexpression is associated with reduced YAP phosphorylation in breast cancer cell spheres and human tumors

Apart from the pro-apoptotic nuclear activities, MST1/MST2 function as core Hippo pathway kinases [16,17]. To test if the FGFR4-dependent suppression of MST1/2 activation alters this canonical Hippo tumor suppressor axis, we analyzed YAP S127 phosphorylation and localization in 2D and 3D cultures of MDA-MB-453 shScr and shFGFR4 cells. In 2D, FGFR4 silencing did not consistently alter YAP S127 phosphorylation or nuclear localization, which both varied depending on the level of serum and cell confluency (Fig. S4A-E). Although the nuclear/cytoplasmic intensity ratio remained essentially unaltered (Fig. S4F), FGFR4 silencing increased the inactive YAP pS127 relative to total YAP in the 3D spheres (Fig. 5A). Moreover, dispersed YAP localization in shScr spheres shifted to a pattern of membrane-proximal YAP after FGFR4 silencing, coincident with increasingly polarized cell architecture and epithelial-like junctions (Fig. 5B).

**Figure 5.**
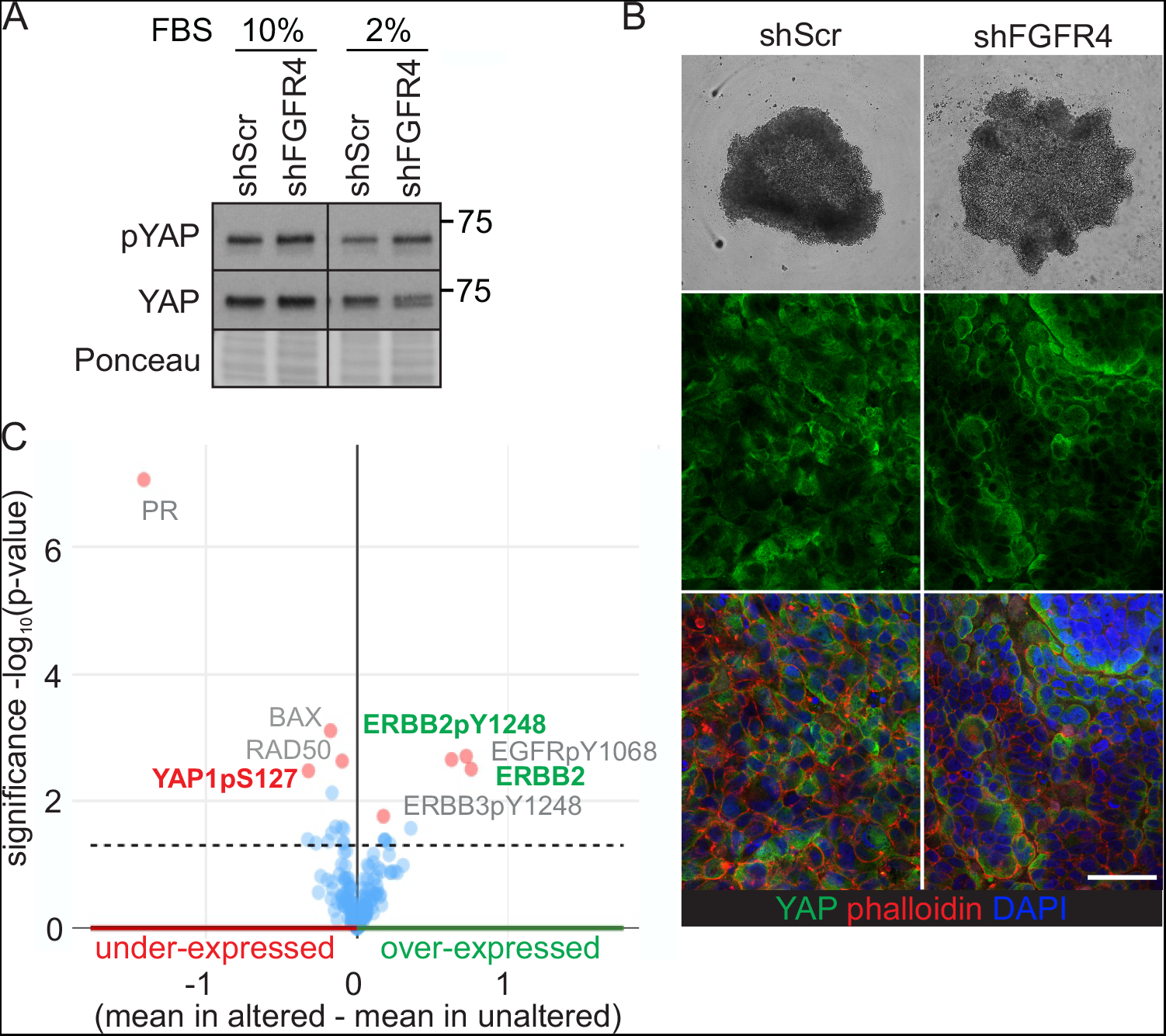
Suppression of YAP phosphorylation in FGFR4 expressing cell spheres concurs with YAP deregulation in breast cancer tumors. **(A)** shScr and shFGFR4 MDA-MB-453 spheres were subjected to immunoblotting for pS127 and total YAP; Ponceau staining as loading control. **(B)** Light micrographs, and immunofluorescence for YAP and filamentous actin (phalloidin) of the MDA-MB-453 cell spheres. Scale bar, 50 μm. **(C)** (Phospho)protein changes in TCGA RPPA data [4] associated with FGFR4 upregulation in breast cancer, visualized using cBioPortal [38,39]. The four most significantly up- and downregulated proteins are highlighted (red dots); ERBB2, alternative name of HER2; PR, progesterone receptor.

To obtain a further unbiased view of the identified functional link between FGFR4 and Hippo pathway in human breast cancer, we systematically analyzed (phospho)protein alterations upon FGFR4 overexpression using TCGA reverse phase protein array (RPPA) data. The most significant alteration associated with FGFR4 overexpression was the decreased progesterone receptor expression (Fig. 5C). Consistent with the co-expression with HER2 mRNA, FGFR4 overexpression likewise correlated with increased total and pY1248 HER2 (ERBB2), as well as EGFR pY1068 and ERBB3 pY1248 in conjunction with adverse patient outcome (Fig. 5C, S4G; TCGA [4]). Significantly, YAP pS127 was downregulated in the FGFR4 overexpressing tumors (Fig. 5C), supporting the FGFR4 function in promoting aggressive properties in human HER2^+^ breast cancer by down-regulation of both MST1/2 and YAP in 3D microenvironment.

## Discussion

Apoptosis evasion is one of the classical hallmarks of cancer. In this study, we identified the apoptosis-promoting MST1/2, together with the other Hippo pathway proteins CSNK1D and PRKCI, among the top five FGFR4 substrates based on Z-score in an unbiased *in vitro* screen. We further provide evidence suggesting that the FGFR4-dependent MST1 Y433 phosphorylation inhibited MST1/2 activity. In the endogenous FGFR4+/HER2^+^ breast cancer cell model, this inhibition was essential to counteract the induction of MST1/2-dependent, stress-associated apoptosis. This suggests that by the identified mechanism, FGFR4 increases the cancer cell survival potential in these breast tumors, where proliferation is driven by HER/ERBB as well as FGFR4 signaling [4,5]. Additionally supporting this conclusion, FGFR4 overexpression was associated with poor outcome of the HER2^+^ breast cancer patients.

MST1 and MST2 (orthologs of *Drosophila* Hippo) are the core kinases of the Hippo signaling pathway. However, the complex and highly context-dependent regulatory mechanisms of mammalian Hippo pathway in health and disease still remain incompletely understood. What is known is that the MST1/2 are activated via homo- or heterodimerization followed by trans-phosphorylation at T183/180 [17,44]. Upon apoptosis, caspase-3-mediated cleavage removes the C-terminal regulatory and nuclear export signals of MST1/2, triggering nuclear translocation of the active N-terminal serine/threonine kinase [18,19]. Although MST1 overexpression has previously been implicated in histone phosphorylation and chromatin condensation [19,45], the contribution of endogenous MST1/2 to apoptosis, downstream of caspase-3 activation, has remained unclear [17]. In this study, we show that in the FGFR4^+^/HER2^+^ breast cancer cells, particularly under the tumor mimicking conditions of 3D spheres in low serum, FGFR4 silencing induced MST1/2 T183/180 autophosphorylation in conjunction with prominent MST1 cleavage (MST1/N), as wells as MST1 nuclear localization without any additional stimuli. This was coupled with significant apoptosis induction via active MST1/2, since the increased apoptosis was rescued back to low control levels by double-knockdown of FGFR4 with MST1 or MST2. In the presence of endogenous FGFR4, however, MST1 or MST2 silencing only did not markedly alter apoptosis, suggesting that these breast cancer cells have developed dependency of the herein identified FGFR4-mediated suppression of this stress-induced apoptosis machinery.

Previously, both positive and negative MST1/2 regulation by other kinases have been reported [27,46–49]. In neuronal cells, the non-receptor tyrosine kinase c-Abl can phosphorylate MST1 at Y433 (conserved site in mammals, absent in MST2) leading to MST1 stabilization and activation in conjunction with increased cell death [50], which is an opposite outcome from the herein identified FGFR4-mediated inhibition of MST1/2 and apoptosis. Moreover, c-Abl-mediated phosphorylation of Y81 on MST2 (species-conserved site, absent in MST1) enhances MST2 activation [51]. The specific aspects of MST2 tyrosine phosphorylation, (hetero)dimerization and activity or cleavage regulation in different cells and conditions will still remain interesting subjects for future studies. Our results also raise new questions regarding possible interrelated functions of Abl and FGFR4 in MST1/2 regulation in cancer cells, since Abl1 and Abl2 also scored as hits in our FGFR4 substrate screen (data not shown). Of note, the other herein identified unique MST1 phosphosite pS410 was detected only in the absence of FGFR4 activity and pY433, raising the possibility of mutually regulated phosphorylation of these MST1 residues. The two MST1 phosphosites detected independently of pY433 and FGFR4 have also been characterized previously; MST1 S320 as a phosphorylation site for anti-apoptotic CK2 [52], and T177 as a MST1 activity-dependent site [53]. Therefore, the specific net effects of all the MST1 phosphosites on its activity regulation, including context-dependent contributions by kinases other than FGFR4, will be of interest. Nevertheless, our current results show that the active FGFR4 tyrosine kinase phosphorylates and inactivates the proapoptotic MST1/2 serine/threonine kinase in breast cancer cells, thus revealing a novel mechanism of RTK-mediated apoptosis evasion and oncogenic FGFR4 function in cancer.

Apart from the direct pro-apoptotic nuclear functions, MST1/2 act on canonical Hippo signaling, which negatively regulates the oncogenic YAP activity by serine phosphorylation and cytoplasmic retention [17,54]. In cancer, however, the non-canonical regulation of YAP expression, localization and activity frequently prevails in conjunction with the loss of balancing feed-back mechanisms of normal cellular homeostasis [16,54,55]. Therefore, we tested the hypothesis that FGFR4-mediated attenuation of MST1/2 activity could canonically suppress YAP phosphorylation and cytoplasmic retention, thus increasing the nuclear YAP. Indeed, YAP pS127 was increased in FGFR4 knockdown 3D cell spheres, and decreased in the FGFR4-overexpressing human HER2^+^ breast cancers in TCGA RPPA data. The multidimensional microenvironment seems crucial for this, as in 2D cell cultures the FGFR4-mediated MST1/2 suppression did not correlate with YAP S127 phosphorylation or nuclear translocation. This is consistent with accumulating evidence indicating that factors such as cell polarity, junctional complexes, and cytoskeletal signals are important in YAP regulation even independent of MST1/2 [15,48]. An interesting further aspect for these cellular regulatory mechanisms comes from the findings that *FGFR1, −2,* and *−4* are direct transcriptional targets of YAP, and a feed-forward loop has been described between YAP and FGFR signaling at least in cholangiocarcinoma and ovarian cancer [56,57].

In conclusion, the identified oncogenic FGFR4 activity explains mechanistically how RTKs such as FGFR4 can confer aggressiveness to certain cancer cells via apoptosis resistance. Since FGFR4 and its kinase activity are druggable targets, these results raise tempting questions, whether inhibition of overexpressed FGFR4 in HER2^+^ breast cancer would release the intrinsic stress-induced MST1/2-mediated apoptotic machinery. And if so, would it offer increase in efficacy in combination with HER2 targeting in this cancer subset, or additionally also in other FGFR4 overexpressing cancer types? Nonetheless, our current identification of MST1/2 as direct substrates for FGFR4, and FGFR4-dependent inhibition of apoptosis and Hippo pathway, highlights interesting avenues for FGFR4 targeting in anti-cancer therapies.

## Acknowledgements

We thank Anastasiya Chernenko for excellent technical assistance, Dr. Jin Cheng (Motiff Cancer Center, Florida, USA) for Flag-tagged MST1 and MST2 plasmids, Julia Casado for initial TCGA expression analysis, and Biomedicum Imaging Unit, University of Helsinki for imaging facilities. This work was funded by the Karolinska Institutet, KI Strategic Research Program in Cancer (StratCan-KICancer), Swedish Cancer Society (Cancerfonden), Swedish Research Council (Vetenskapsrådet), the University of Helsinki, Academy of Finland, Sigrid Jusélius Foundation, Magnus Ehrnrooth Foundation, Medicinska understödsföreningen Liv och hälsa, and the The Finnish Society of Sciences and Letters.

## Supplemental Figures and legends

**Figure S1.**
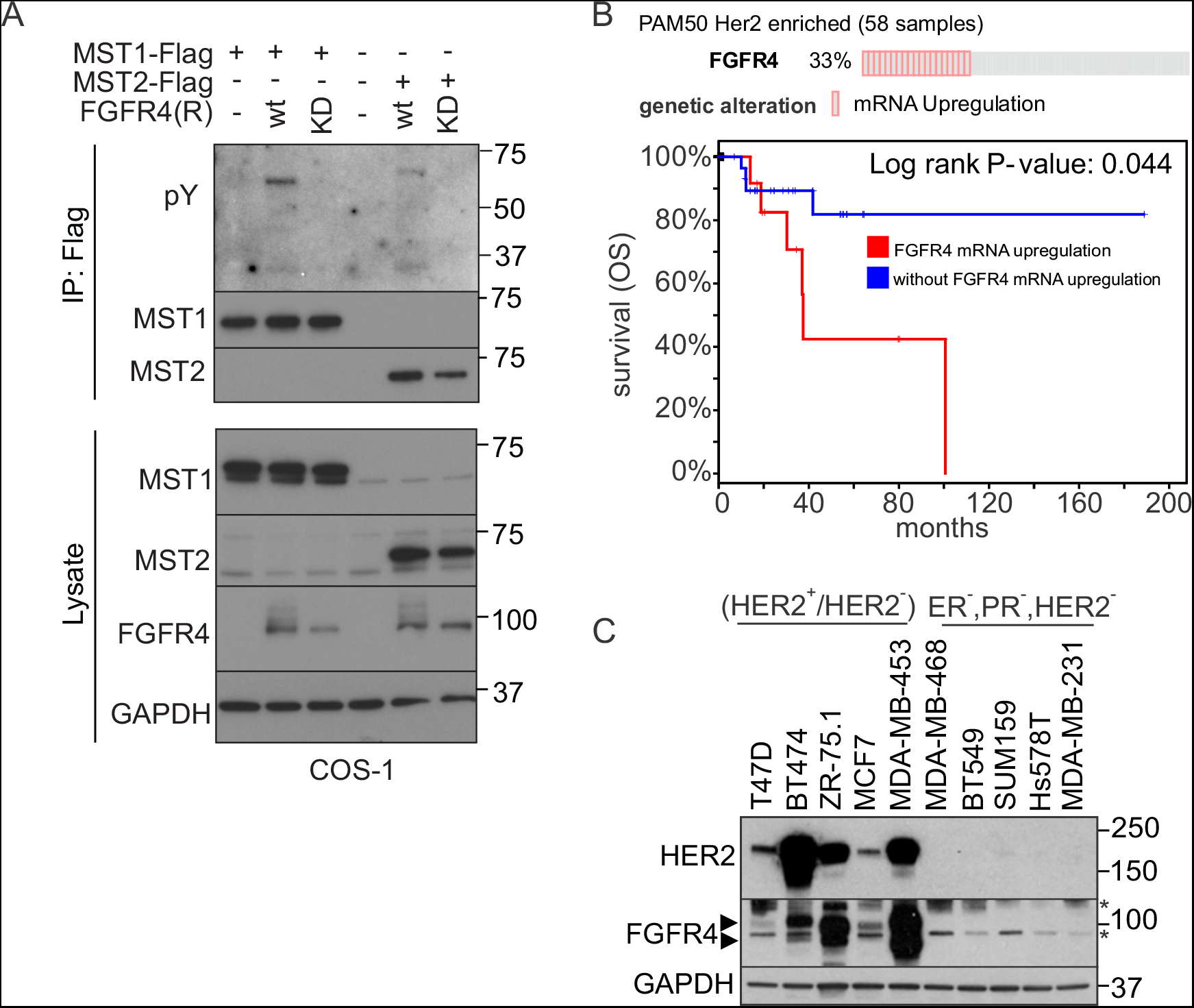
Tyrosine phosphorylation of MST1/2 in COS-1 cells co-transfected with FGFR4. FGFR4 survival association in breast cancer TCGA data and expression in cell lines. **A)** Flag-tagged MST1/2 were immunoprecipitated from COS-1 cells after transfection of MST1 and MST2 alone or in combination with FGFR4 R388 (R) kinase (wt) or kinase-dead (KD) variants, and detected by immunoblotting. For phosphorylation site identification, the urea-eluted MST1 immunoprecipitates from COS-1 cells co-transfected with FGFR4 (R)-wt or FGFR4 (R)-KD were trypsin digested and subjected to phosphopeptide enrichment prior to LC-MS/MS analysis. **B)** Kaplan-Meier survival curve of patients with HER2 positive breast cancer (TCGA, PAM50 classification: HER2 enriched, cBioPortal for Cancer Genomics) visualizes the correlation between FGFR4 upregulation and poor overall survival (OS). **C)** Immunoblotting of HER2 and FGFR4 in a set of luminal and triple-negative breast (ER^−^, PR^−^, HER2^−^) cancer cell lines. Long exposures of HER2 and FGFR4 corresponding to immunoblot in Fig. 2A. Arrow heads denote FGFR4, asterisks mark unspecific bands.

**Figure S2.**
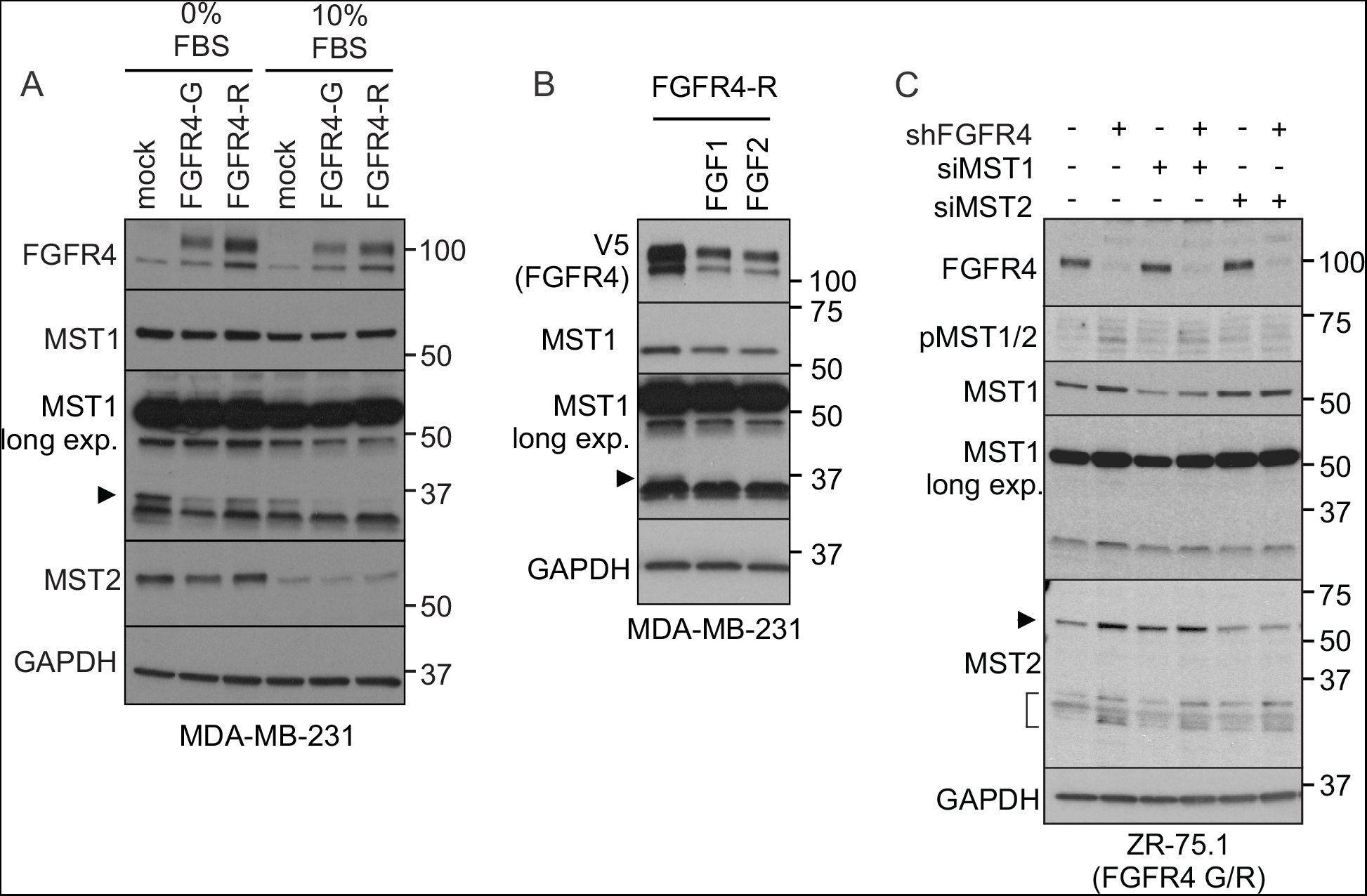
Reduced MST1 cleavage in FGFR4 overexpressing MDA-MB-231 cells, and activation of MST1/2 in ZR-75.1 cells upon FGFR4 silencing. **A, B)** MDA-MB-231 cells transfected and selected for stable overexpression of FGFR4-(R) or -(G) were cultured **(A)** without serum or with 10% FBS, or **(B)** stimulated with 10 ng/ml of FGF1 or FGF2 for 16 hours in serum-free medium, and subjected to immunoblotting. Arrow head points to a cleaved MST1 (~37 kDa). (**C)** ZR-75.1 cells were transduced with shRNAs followed by transfection with siRNAs as indicated, and subjected to immunoblotting for pT183/180 MST1/2, MST1 and MST2 (arrow head points to a full-length, bracket to the cleaved protein).

**Figure S3.**
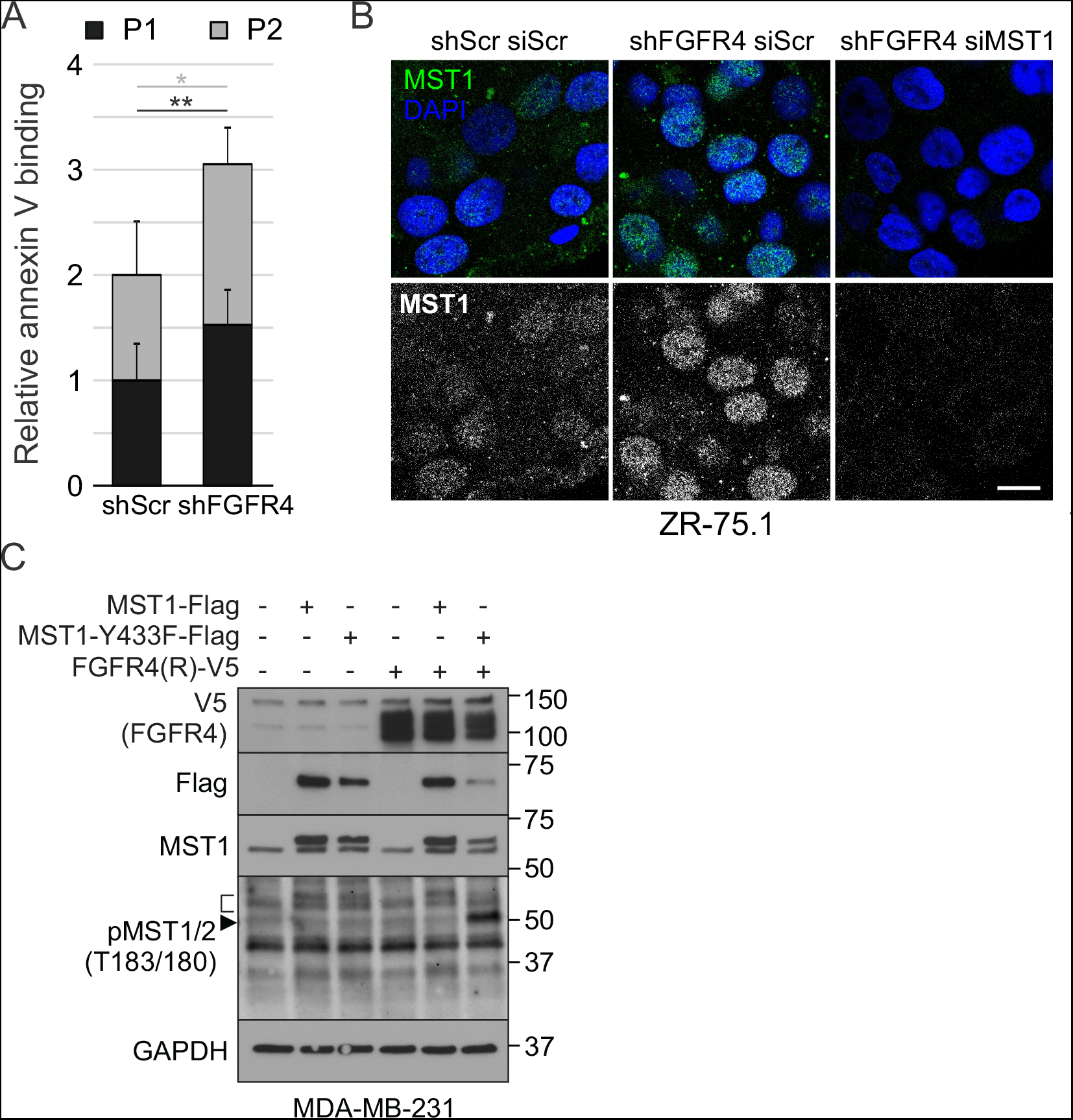
FGFR4 silencing increases apoptosis in breast cancer cells. MST1 localizes to nucleus upon FGFR4 silencing in ZR-75.1 cells. **A)** Flow cytometry analysis and quantification of relative annexin V binding in MDA-MB-453 cells in populations P1 and P2; * P<0.05; ** P<0.01. **B)** ZR-75.1 cell spheres transduced with shScr or shFGFR4 shRNAs and transfected with indicated siRNAs were analyzed for MST1 localization by immunofluorescence. **(C)** MDA-MB-231 cells (co-)transfected with wild-type or phosphosite mutant MST1-Y433F and FGFR4 (R) were subjected to immunoblotting as indicated. Bracket and arrow head indicate the activated pT183 MST1 fragments.

**Figure S4.**
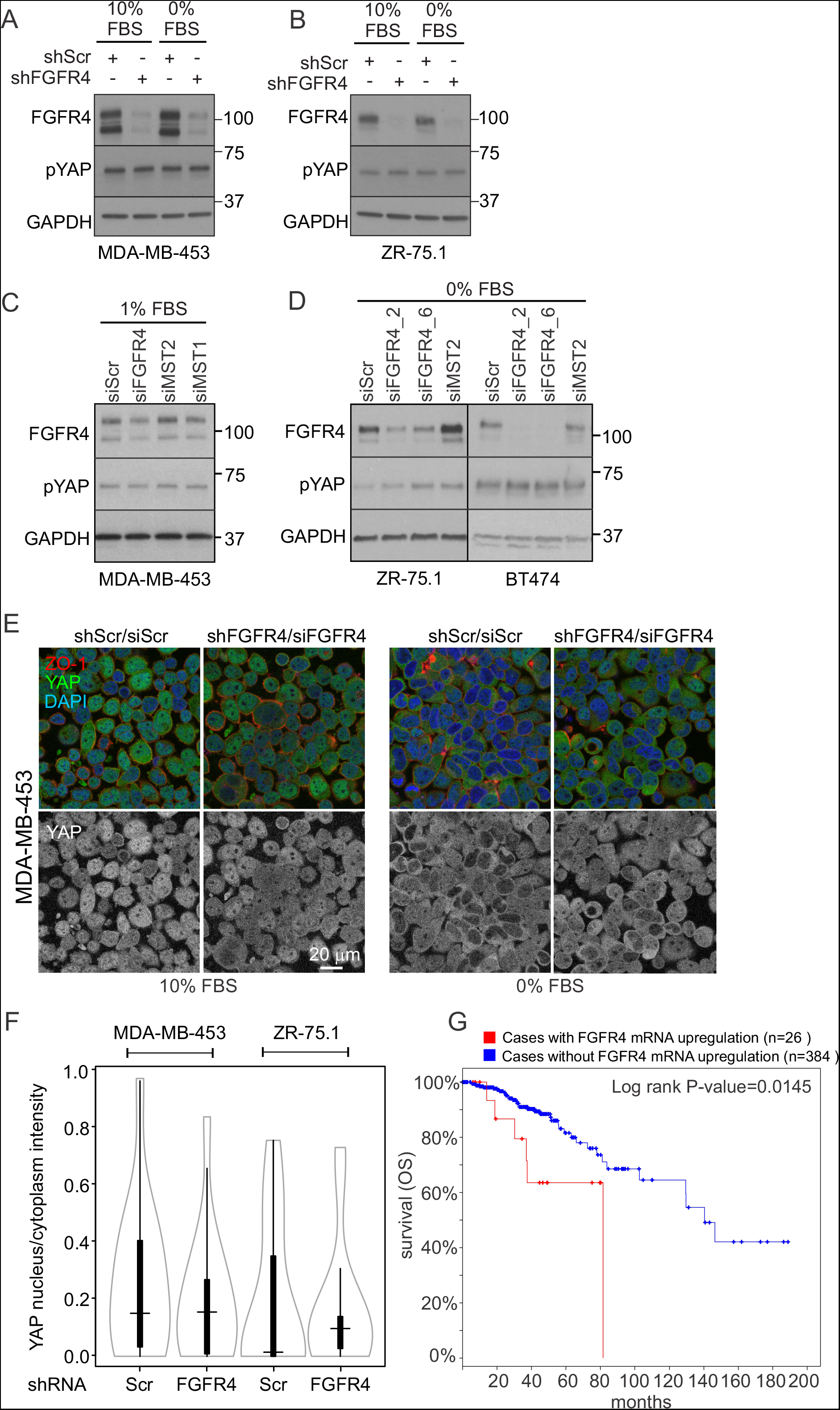
YAP nuclear localization remains unaltered in experimental models of high endogenous FGFR4 expression in breast cancer cells. **(A)** MDA-MB-453 and **(B)** ZR-75.1 cells were transduced with indicated shRNAs, cultured in serum-free (0% FBS) or 10% FBS containing medium, and subjected to immunoblotting for FGFR4, pS127 YAP, and GAPDH as a loading control. **(C)** MDA-MB-453 cells transfected with indicated siRNAs were cultured in medium with 1% FBS, **(D)** ZR-75.1 and BT474 cells transfected with indicated siRNAs were cultured in serum-free medium, and subjected to immunoblotting for FGFR4, pS127 YAP, and GAPDH as a loading control. (**E)** Immunofluorescence staining of YAP and ZO-1 (zona occludens-1) in MDA-MB-453 cells transfected with indicated siRNAs, and cultured in 2D with (10%) or without FBS (0%). (**F)** Quantification of YAP nucleus/cytoplasma intensity ratio in MDA-MB-453 and ZR-75.1 spheres formed from cells transduced with shScr or shFGFR4 shRNAs. **(G)** Kaplan-Meier survival curve of patients included in the RPPA of the TCGA Nature 2012 cohort (n=410 tumors (Cancer Genome Atlas Network 2012), cBioPortal for Cancer Genomics (Cerami et al., 2012, Gao et al., 2013)) visualizes the correlation between FGFR4 alterations and poor overall survival (OS) in breast cancer.

